# A powerful and versatile new fixation protocol for immunohistology and *in situ* hybridization that preserves delicate tissues in planaria

**DOI:** 10.1101/2021.11.01.466817

**Authors:** Carlos Guerrero-Hernández, Viraj Doddihal, Frederick G. Mann, Alejandro Sánchez Alvarado

## Abstract

Whole-mount *in situ* hybridization (WISH) is a powerful and widely used technique to visualize the expression pattern of genes in different biological systems. Here we describe a new protocol for ISH and immunostaining in the planarian *Schmidtea mediterranea*. The new Nitric Acid/Formic Acid (NAFA) protocol is compatible with both assays and prevents degradation of the epidermis or blastema. Instead of proteinase K digestion, formic acid treatment is used to permeabilize tissues and preserve antigen epitopes. We show that the NAFA protocol successfully permits development of chromogenic and fluorescent signals *in situ*, while preserving the anatomy of the animal. Further, the immunostaining of different proteins was compatible with the NAFA protocol following fluorescent *in situ* hybridization. Finally, we demonstrate with high resolution confocal imaging that the regeneration blastema is preserved when using the new method. This new NAFA protocol will be a valuable technique to study the process of wounding response and regeneration.

## Introduction

The freshwater planarian *S. mediterranea* can regrow a complete animal from a body fragment that is less than 1% of its original size^1^. This remarkable capacity for regeneration has attracted the attention of generations of biologists and its study has required the development of methods to detect, measure, and visualize the cells and molecules underpinning this process. Planarians respond to wounds by the spreading of the ciliated epidermis over the surface of the damaged area. This is followed by proliferation of the stem cells (called neoblasts) and the generation of a blastema, an undifferentiated structure produced at the wound site that is necessary for constructing new tissue and restoring the missing body parts^2^. Understanding how genes function in the epidermis and blastema to heal wounds and restore lost tissue is essential to understanding regeneration. Yet, the existing methods for studying gene expression in the context of a whole planarian are limited. Transgenesis is not currently possible in *S. mediterranea* and only a few antibodies are available. As a result, RNA *in situ* hybridization (ISH) has been a primary tool for studying the biology of planarian stem cells and regeneration^3,4^. This method has revealed genes that control the asymmetric division^5^ and specification of stem cells^6,7^, as well as cues from differentiated cells that modulate stem cell proliferation^8,9^, relay information about polarity^10,11^, and help define position in the body^12^.

However, current ISH protocols have several shortcomings. Penetration of probes into tissue for whole-mount *in situ* hybridization (WISH) is difficult to achieve. As such, permeability is increased through tissue digestion with proteinase K, and through aggressive treatment with N-acetyl cysteine (NAC)^3,4^. These harsh treatments can damage or destroy fragile tissues, and often result in the shredding of both the epidermis and the regeneration blastema. Even so, probe penetration into larger animals is limited. Moreover, immunological assays are generally weak on samples prepared by this protocol, likely because proteinase digestion disrupts target epitopes. Other protocols have been developed for fixing whole planarians that preserve the gross anatomical structures and perform well in immunological assays^13-15^. Unfortunately, that methodology is not compatible with ISH. An ideal method would preserve delicate tissues and permit the simultaneous analysis of RNA and protein expression patterns.

Here, we present a new protocol for ISH and immunofluorescence in planarians. We have combined approaches from several fixation strategies into a nitric acid / formic acid strategy for sample preparation^13-15^ that better preserves the delicate epidermis and blastema than previous methods do. Our new protocol increases probe penetration, resulting in stronger signals. As a result, samples do not require proteinase digestion, and therefore have greatly increased compatibility with immunological assays. Our protocol allows delicate tissues to be studied more readily via ISH and immunofluorescence, which will lead to a better understanding of regeneration.

## EXPERIMENTAL PROCEDURES

### Materials

All solutions should be made in Milli-Q water

**Nitric acid, 70% (HNO_3_)** Sigma-Aldrich

**Magnesium sulfate (MgSO_4_)** Sigma-Aldrich

**EGTA** Sigma-Aldrich

**HEPES** Sigma-Aldrich

**Formic acid** Sigma-Aldrich

**Acetic acid** Sigma-Aldrich

**Lactic acid** Fisher Chemical

**Formaldehyde solution 36.5-38%** Sigma-Aldrich

**Tween 20 (Polyethylene glycol sorbitan monolaurate)** Sigma-Aldrich

**Triton X-100 (Polyethylene glycol p-(1,1,3,3-tetramethylbutyl)-phenyl ether)** Sigma-Aldrich

**Hydrogen peroxide 30% (H_2_O_2_)** Sigma-Aldrich

**Paraformaldehyde 16%** Electron Microscopy Science

**Methanol** Fisher Chemical

**PBS1X (Standard buffer solution):** 137 mM NaCl, 2.7 mM KCl, 10 mM Na_2_HPO_4_, 2 mM KH_2_PO_4_, in H_2_O, final pH 7.4.

**Formamide (Deionized)** Millipore

### NA Solution

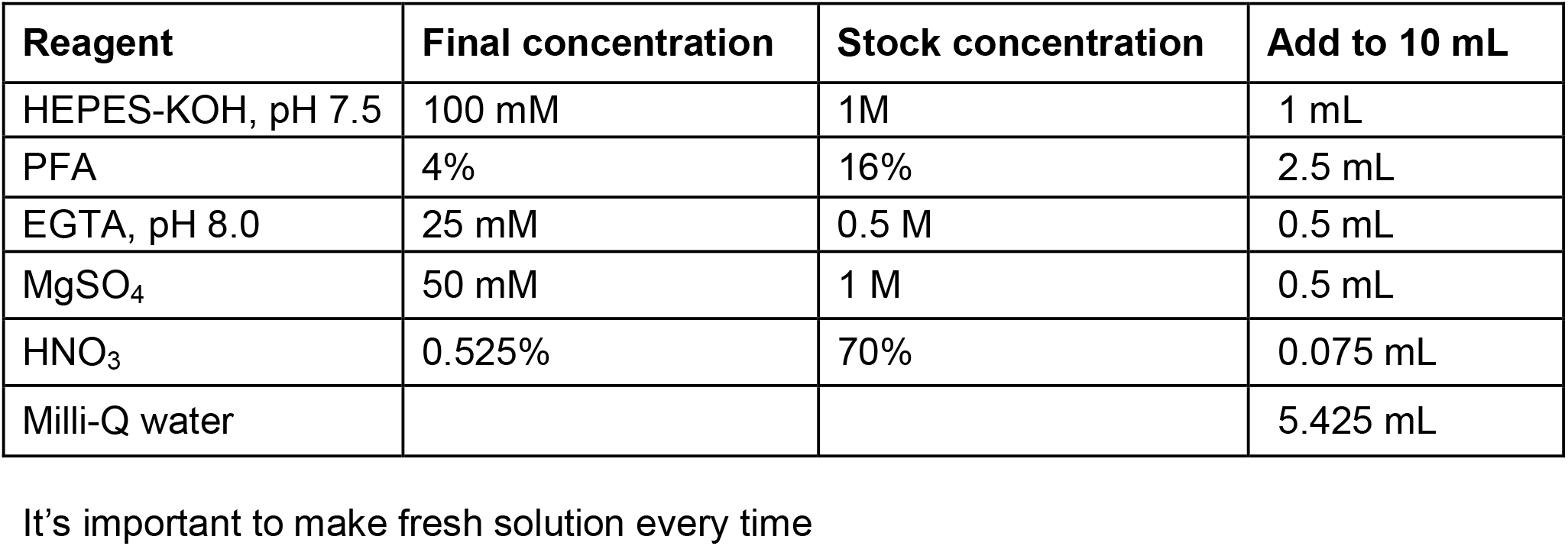

| **Reagent** | **Final concentration** | **Stock concentration** | **Add to 10 mL** |
| --- | --- | --- | --- |
| HEPES-KOH, pH 7.5 | 100 mM | 1M | 1 mL |
| PFA | 4% | 16% | 2.5 mL |
| EGTA, pH 8.0 | 25 mM | 0.5 M | 0.5 mL |
| MgSO<sub>4</sub> | 50 mM | 1 M | 0.5 mL |
| HNO<sub>3</sub> | 0.525% | 70% | 0.075 mL |
| Milli-Q water |  |  | 5.425 mL |

It's important to make fresh solution every time

### FA Solution

| **Reagent** | **Final concentration** | **Stock concentration** | **Add to 10 mL** |
| --- | --- | --- | --- |
| HEPES-KOH, pH 7.5 | 100 mM | 1 M | 1 mL |
| PFA | 4% | 16% | 2.5 mL |
| EGTA, pH 8.0 | 25 mM | 0.5 M | 0.5 mL |
| Formic acid | 4.80% | 96% | 0.5 mL |
| Milli-Q water |  |  | 5.5 mL |

It's important to make fresh solution every time

### Wash Hybe – 0.5 % Tween-20

| Reagent | Final concentration | Add to 500 mL |
| --- | --- | --- |
| Formamide | 50% | 250 mL |
| 10 % Tween-20 | 0.50% | 25 mL |
| 20 X SSC | 5 X | 125 mL |
| 50 X Denhardts | 1 X | 10 mL |
| Milli-Q water |  | Up to 500 mL |

### Pre-Hybe

| Reagent | Final concentration | Add to 500 mL |
| --- | --- | --- |
| Formamide | 50% | 250 mL |
| 10 % Tween-20 | 1% | 50 mL |
| 20 X SSC | 5 X | 125 mL |
| 50 X Denhardts | 1 X | 10 mL |
| Heparin | 100 mg | 0.05 g |
| DTT | 50 mM | 3.85 g |
| Torula Yeast RNA | 1 mg | 0.5 g |
| Milli-Q water |  | Up to 500 mL |

Store at -20°C.

### Hybridization buffer (Hybe)

| Reagent | Final concentration | Add to 500 mL |
| --- | --- | --- |
| Formamide | 50% | 250 mL |
| 10 % Tween-20 | 1% | 50 mL |
| 20 X SSC | 5 X | 125 mL |
| 50 X Denhardts | 1 X | 10 mL |
| Heparin | 100 mg | 0.05 g |
| DTT | 50 mM | 3.85 g |
| Torula Yeast RNA | 1 mg | 0.5 g |
| Dextran Sulfate | 5% | 25 g |
| Milli-Q water |  | Up to 500 mL |

Store at -20°C.

### Nitric acid / formic acid (NAFA) fixation protocol

Note: All the steps are carried out with animals being nutated/rocked at room temperature unless stated otherwise.

1. Transfer planarians starved for at least for one week to either 1.5 mL tubes or 15 mL tubes for processing up to 20 or 100 animals, respectively.
2. Replace planarian water with NA solution for 1-2 minutes. During this treatment agitate animals vigorously by inverting the tubes a few times. NA solution has nitric acid and magnesium sulfate which helps anesthetize (relax) and euthanize the animals prior to fixation. Note: This treatment should not go beyond 5 minutes. NA solution: 0.5% nitric acid, 4% paraformaldehyde, 50mM magnesium sulfate, and 25mM EGTA (pH 8.0) in 100 mM HEPES (pH 7.5).
3. Replace the NA solution with FA solution and incubate the animals in this solution for 40 minutes to 1 hour. Fix solution: 4% paraformaldehyde, 25mM EGTA (pH 8.0), and 4.8% formic acid in 100mM HEPES (pH 7.5)
4. Remove the FA solution and wash twice in 1X PBS for 10 minutes each.
5. Following the 1X PBS washes, wash animals in 50% methanol in 1X PBS for 10 minutes.
6. Replace the 50% methanol in 1X PBS with 100% methanol and incubate for 10 minutes to allow thorough dehydration.
7. Replace the solution with fresh 100% methanol and store in −20°C for at least one hour or until ready to use.
8. When ready to use the fixed specimens, replace the 100% methanol with 50% methanol in 1X PBS for 10 minutes.
9. Once completed, replace the 50% methanol with 1X PBS for 10 minutes.
10. Bleach animals under direct light in formamide bleach solution for 2 hours. Formamide bleach solution: 1% formamide, 6% hydrogen peroxide in PBSTx (0.3% - 0.5% Triton).
11. Rinse the animals twice for 10 minutes each in PBSTx (0.3% - 0.5% Triton).

Following this step, directly proceed to either *in situ* hybridization or immunostaining protocols. There is no proteinase K treatment.

### *in situ* hybridizations

12. Replace PBSTx (0.3% - 0.5% Triton) with 1:1 (PBSTx:PreHybe) solution for 5 minutes.
13. Incubate animals in Pre-Hybe solution for 2 hours at 56°C.
14. Replace Pre-Hybe with riboprobe mix (riboprobe(s) in hybridization buffer) for ≥16 hours at 56°C. Riboprobes are generally used at 1:1000 dilution and can be heat denatured in hybridization buffer at 70°C for 3 minutes prior to use.
15. Carry out post hybridization washes at 56°C
  a. Wash with Wash Hybe two times for 30 minutes each.
  b. Wash with 1:1 mix of Wash Hybe:2X SSC (+ 0.1% Tween or Triton) two times for 30 minutes each.
  c. Wash three times with 2X SSC (+ 0.1% Tween or Triton) for 20 minutes each.
  d. Wash three times with 0.2X SSC (+ 0.1% Tween or Triton) for 20 minutes each
16. Once post-hybridization washes are completed, wash animals three times with MABT, for 10 minutes each at room temperature.
17. Block with 5% filtered horse serum + 0.5% filtered Roche Western Blocking Reagent (RWBR) in MABT for 1-2 hours.
18. Incubate the samples overnight at room temperature, with appropriate antibody, diluted in blocking solution. We regularly use:
  a. Anti-DIG-AP at 1:3000 dilution (Roche – Cat. No. 11093274910)
  b. Anti-DIG-POD at 1:1000 dilution (Roche - Cat. No. 11207733910)
  c. Anti-Fluorescein-POD at 1:3000 dilution (Jackson Laboratories – Cat. No. AB_2314402)
19. Wash animals 6 times in MABT for 20 minutes each.

### Tyramide-based fluorescent signal development

20. For developing fluorescent signal, preincubate the samples with tyramide in borate buffer for 15 minutes.
21. Carry out the tyramide reaction by adding 0.006% hydrogen peroxide in borate buffer for 45 minutes.
22. Stop the reaction by washing out the samples twice in PBSTw (0.3% Tween) for 10 minutes each.
23. If developing a second probe –
  a. Kill the peroxidase activity by treating with 200mM sodium azide in PBSTw (0.3% Tween) for at least 1 hour.
  b. Wash 6 times with PBSTw (0.3% Tween) for 20 minutes each.
  c. Wash 4 times with MABT for 10 minutes each.
  d. Return to step 18 to develop the second probe.
24. If performing immunostaining skip to immunostaining protocol.
25. *Optional*: Post-fix the samples in 4% formaldehyde in PBSTx (0.3% Triton) for 20-30 minutes. Note: If performing immunostaining do not post fix at this stage.
26. Clear the samples for 1-2 days in 20% Scale A2 + DABCO.

### NBT/BCIP based colorimetric signal development

20. After washing out the antibody with MABT (step 19), incubate in AP buffer for 1-2 minutes.
21. Replace the AP buffer with the EQ buffer for 10 minutes.
22. Incubate the animals in DEV buffer + NBT/BCIP till the signal is developed to appropriate levels.
23. Wash away NBT/BCIP by rinsing the samples two to three times in 1X PBS.
24. Post fix the samples in 4% formaldehyde in PBSTx (0.3% Triton) for 20 - 30 minutes.
25. Incubate the animals in 100% ethanol to clear the background.
26. Rinse the animals with 50% ethanol in 1X PBS for 5 minutes.
27. Wash the samples in 1X PBS for 5-10 minutes or until they sink.
28. Rinse the samples a couple more times in 1X PBS.
29. Clear in 80% glycerol or 75% Scale A2 for 1-2 days.

### Immunostaining

| **Antibody** | **Dilution** | **Source** | **Supplier** | **Catalog Number** |
| --- | --- | --- | --- | --- |
| Anti – H3P | 1:500 – 1:1000 | Rabbit | Abcam | ab32107 |
| Anti – acetylated tubulin | 1:1000 | Rabbit | Cell Signaling | #5335 |
| Anti – acetylated tubulin | 1:1000 | Mouse | Sigma | T7451 |
| 6G10-2C7<br>(muscle<br>antibody) | 1:200 – 1:400 | Mouse | Developmental Studies<br>Hybridoma Bank<br>(DSHB) | 6G10-2C7 |
| Anti-rabbit,<br>Alexa Fluor<br>555 | 1:500 – 1:1000 | Goat | Abcam | 150086 |
| Anti-rabbit,<br>Alexa Fluor<br>647 | 1:500 – 1:1000 | Goat | Abcam | 150083 |
| Anti-mouse,<br>Alexa Fluor<br>555 | 1:500 – 1:1000 | Goat | Abcam | 150118 |
| Anti-mouse,<br>Alexa Fluor<br>647 | 1:500 – 1:1000 | Goat | Abcam | 150119 |

Note: If performing immunostaining on samples after developing *in situ* hybridization signal then start by blocking with 5% goat serum (step 1).

1. 1. Fixed and bleached samples are blocked with 5% goat serum in PBSTw (0.3% - 0.5% Tween) for 1-2 hours.
2. 2. Incubate the samples overnight with primary antibody in blocking solution (5% goat serum in PBSTw (0.3% - 0.5% Tween).
3. 3. Wash the samples six times in PBSTw (0.3% - 0.5% Tween) for 20 minutes each.
4. 4. Incubate overnight with appropriate secondary antibody (1:500 −1: 1000) in blocking solution. Note: DAPI can be added at this step.
5. 5. Wash the samples six times in PBSTw (0.3% - 0.5% Tween) for 20 minutes each.
6. 6. Optional: Post fix the samples in 4% formaldehyde in PBSTx (0.3% Triton) for 20 - 30 minutes.
7. 7. Clear the samples for 1-2 days in 20% Scale A2 + DABCO. Note: We have not observed any significance difference when PBSTx (0.3% −0.5% Triton) is used in place of PBSTw (0.3% - 0.5% Tween).

## Results

We sought to create a new fixation protocol for planarians that would be compatible with both ISH and antibody-based assays while preserving the structural integrity of the animals. We reasoned that combining the acid treatment strategies of a variety of protocols could make the samples compatible with multiple applications^3,13,14,16^. We also included EGTA to inhibit nucleases and preserve RNA integrity during sample preparation^17^. We first determined the extent to which the new combination of acids preserved the samples. We used the integrity of the epidermis as a proxy for tissue preservation, which we visualized by immunostaining cilia with an anti-acetylated tubulin antibody^18^. We tested a <u>N</u>itric <u>A</u>cid / <u>F</u>ormic <u>A</u>cid (NAFA) fixation and compared it against two well established fixation protocols in the field, Nitric Acid (NA)^13^ and N-Acetyl-Cysteine (NAC)^3^. We found that the integrity of the epidermis is well preserved in both the NA and NAFA protocols, whereas noticeable breaches of integrity were detected when the protocol using the mucolytic compound N-acetyl-cysteine was tested (Figure 1). We concluded from these results that the NAFA protocol worked as well as the NA protocol but preserved the sample significantly better than the NAC protocol did. This result prompted us to explore whether the NAFA protocol is also compatible with WISH.

**Fig 1.**
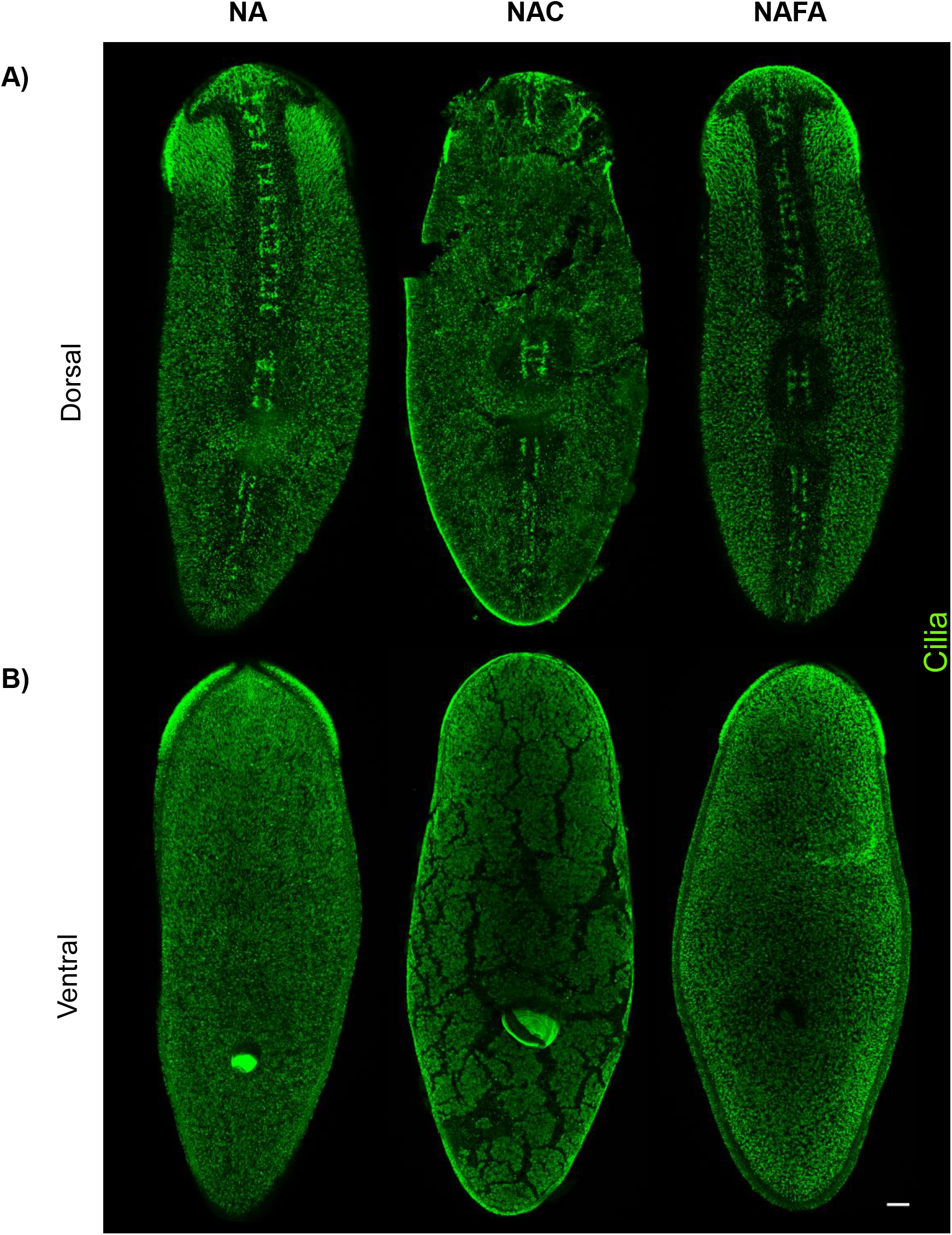
Epidermal integrity is preserved with Nitric acid/Formic acid (NAFA) protocol. Anti-acetylated tubulin staining of cilia on the **(A)** dorsal and **(B)** ventral surfaces. NA: Nitric acid^13^ and NAC: N-acetyl-cysteine^3^. Maximum intensity projection of confocal images, scale bar: 100 μm

Given the success of the immunolabeling, we wished to test whether the NAFA protocol could be used for ISH assays. To ensure the NAFA protocol allows *in situ* probes to penetrate into tissues, we used probes that are known to mark an internal cell population, the neoblasts (*piwi-1*)^19^ and a more external cell population, a subset of the epidermal progenitors (*zpuf-6*). First, we tested whether the expression of *piwi-1* and *zpuf-6*^8^ could be detected via chromogenic WISH (Figure 2). While the NAFA and NAC protocols produced indistinguishable patterns of expression for the two genes, we could not observe any *piwi-1* and *zpuf-6* signal with the NA protocol (Figure 2A and 2B). These experiments also revealed epidermal damage when NAC was used (Figure 2B). To further investigate epidermal integrity and WISH signal, we performed chromogenic WISH for *zpuf-6* using the NAC and NAFA protocols (Figure S1), then sectioned the animals afterwards for histological analysis. The sections revealed that the outermost layer with *zpuf-6+* cells was intact when using the NAFA protocol, but damaged by the NAC protocol (Figure S1A and S1B). Also, we tested whether three different carboxylic acids (formic acid, acetic acid and lactic acid) can be used in the NAFA protocol. We performed chromogenic WISH for *piwi-1, zpuf-6*, in addition to markers of the central nervous system (*pc2*)^20,21^ and gastrovascular system (*porcupine*)^10^. All showed similar expression patterns in both the NAFA and NAC protocols (Figure S2). While all three carboxylic acids can be used to determine gene expression patterns and are effective across multiple transcripts, for further characterization we chose to use the acid with the lowest pKa (Formic acid). In conclusion, the new NAFA protocol both preserves epidermis integrity and can be applied to successfully detect gene expression in different planarian tissues by WISH.

**Fig 2.**
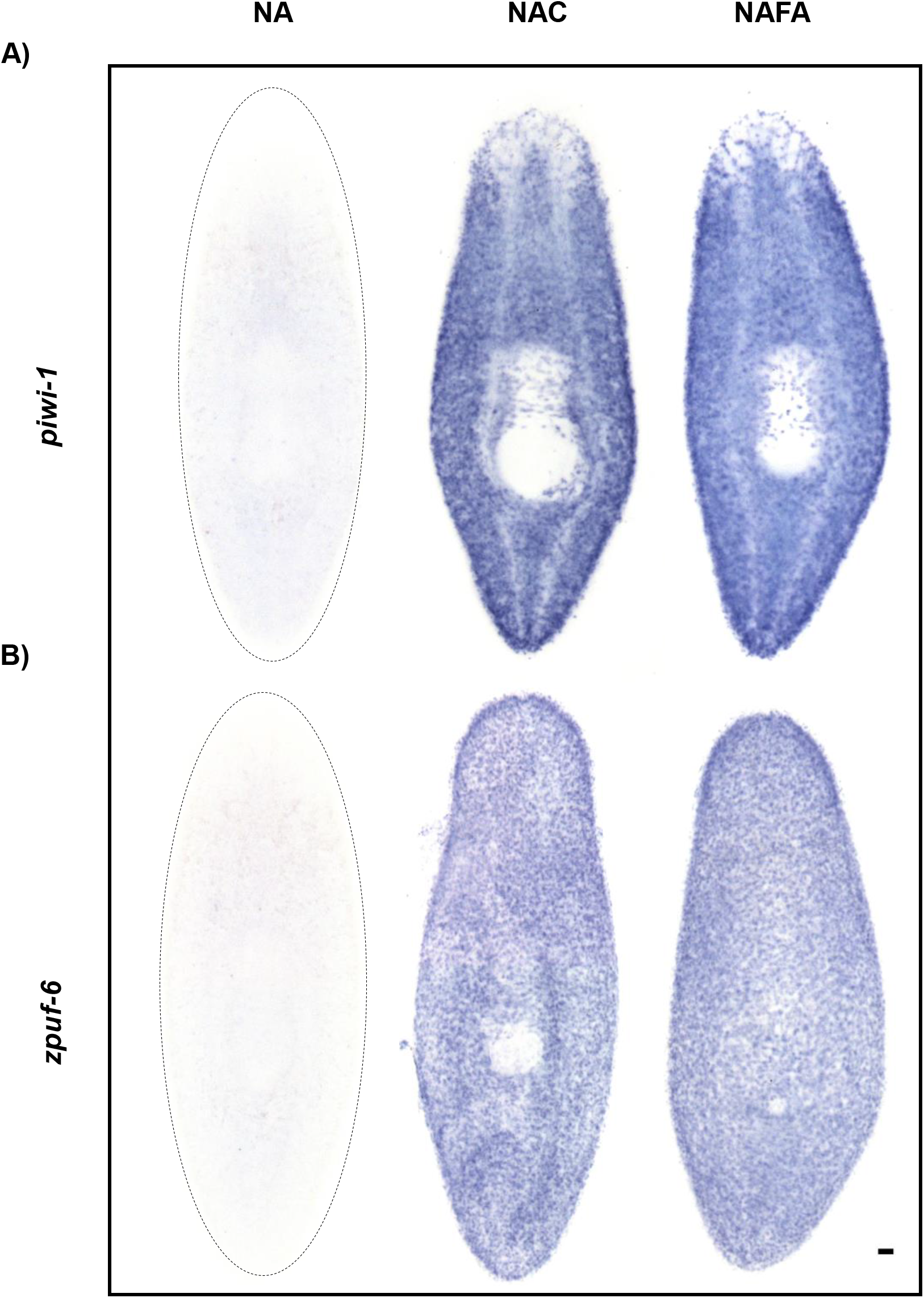
The NAFA protocol is compatible with chromogenic *in situ* hybridization. **(A)** *piwi-1* (neoblasts) and **(B)** *zpuf-6* (epidermal progenitors). Scale bar: 100 μm

Next, we investigated whether we could use the new NAFA protocol in planaria to carry out fluorescent *in situ* hybridization (FISH) in tandem with immunostaining. Confocal microscopy was performed to detect the neoblast and epidermal progenitor markers *piwi-1* and *zpuf-6*, respectively (Figure 3). FISH showed that the tissue specificity of fluorescent detection compared well with that of chromogenic detection of *piwi-1* and *zpuf-6* with both NAC and NAFA protocols (Figure 2 and 3). Furthermore, confocal microscopy showed that the epidermis was damaged with the NAC protocol but was not affected when using the NAFA protocol (Figure 3B). After whole-mount FISH, we immunostained for mitotic cells with an antibody that recognizes the phosphorylated form of Histone H3 (anti-H3P)^22,23^. To our surprise, the anti-H3P antibody showed stronger signal with the NAFA protocol when compared to both NA and NAC protocols (Figure 4A). We also evaluated the compatibility of the three protocols with immunofluorescence by staining the muscle fibers and cilia using the antibodies Smed-6G10^24^ and anti-acetylated tubulin respectively. For both antibodies, the expression patterns were similar with the NA and NAFA protocols, while the signals were much weaker with the NAC protocol (Figure 4B and 4C).

**Fig 3.**
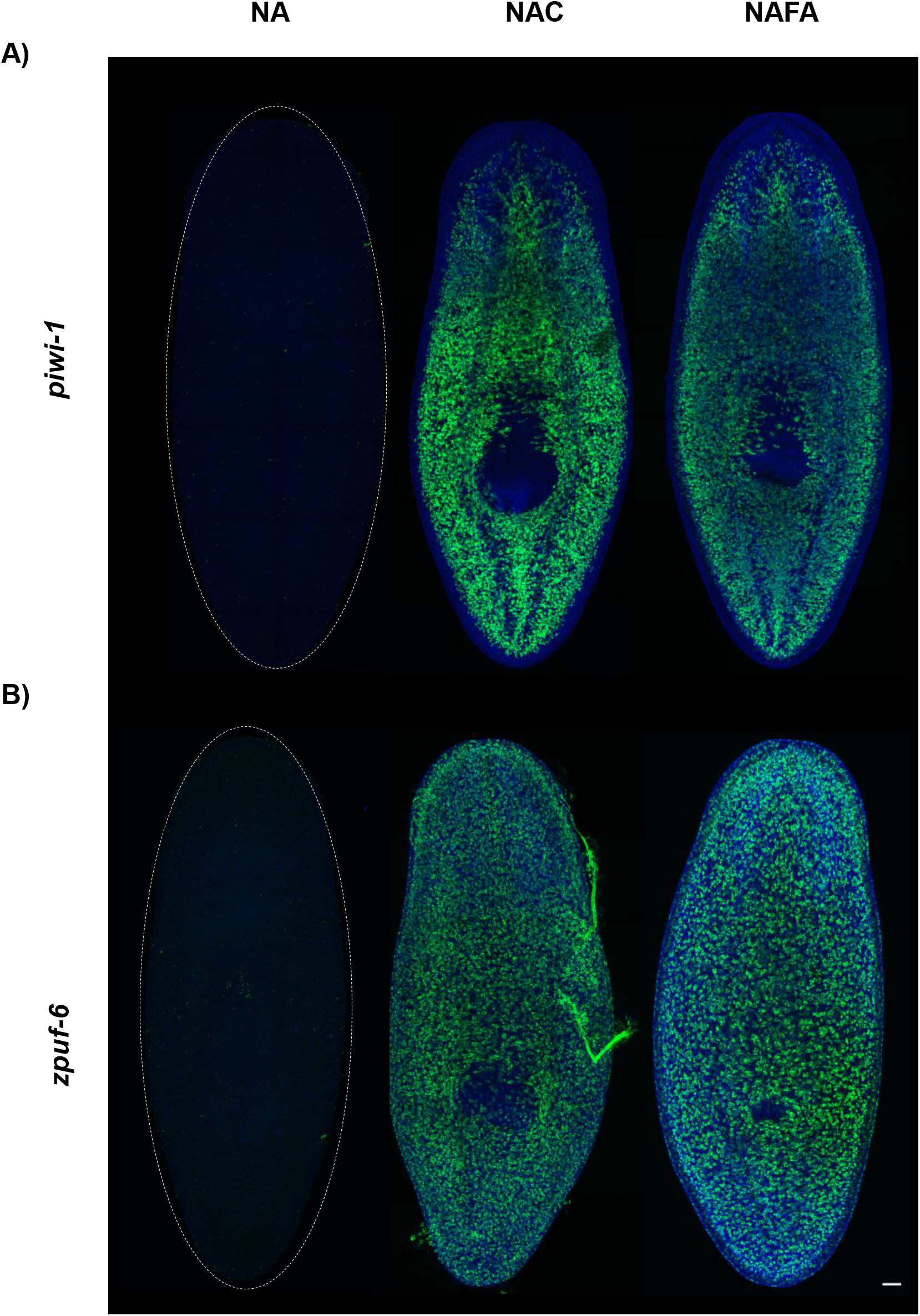
NAFA is compatible with fluorescent *in situ* hybridization. FISH of **(A)** *piwi-1* (neoblasts) and **(B)** *zpuf-6* (epidermal progenitors). Maximum intensity projection of confocal images, scale bar: 100 μm

**Fig 4.**
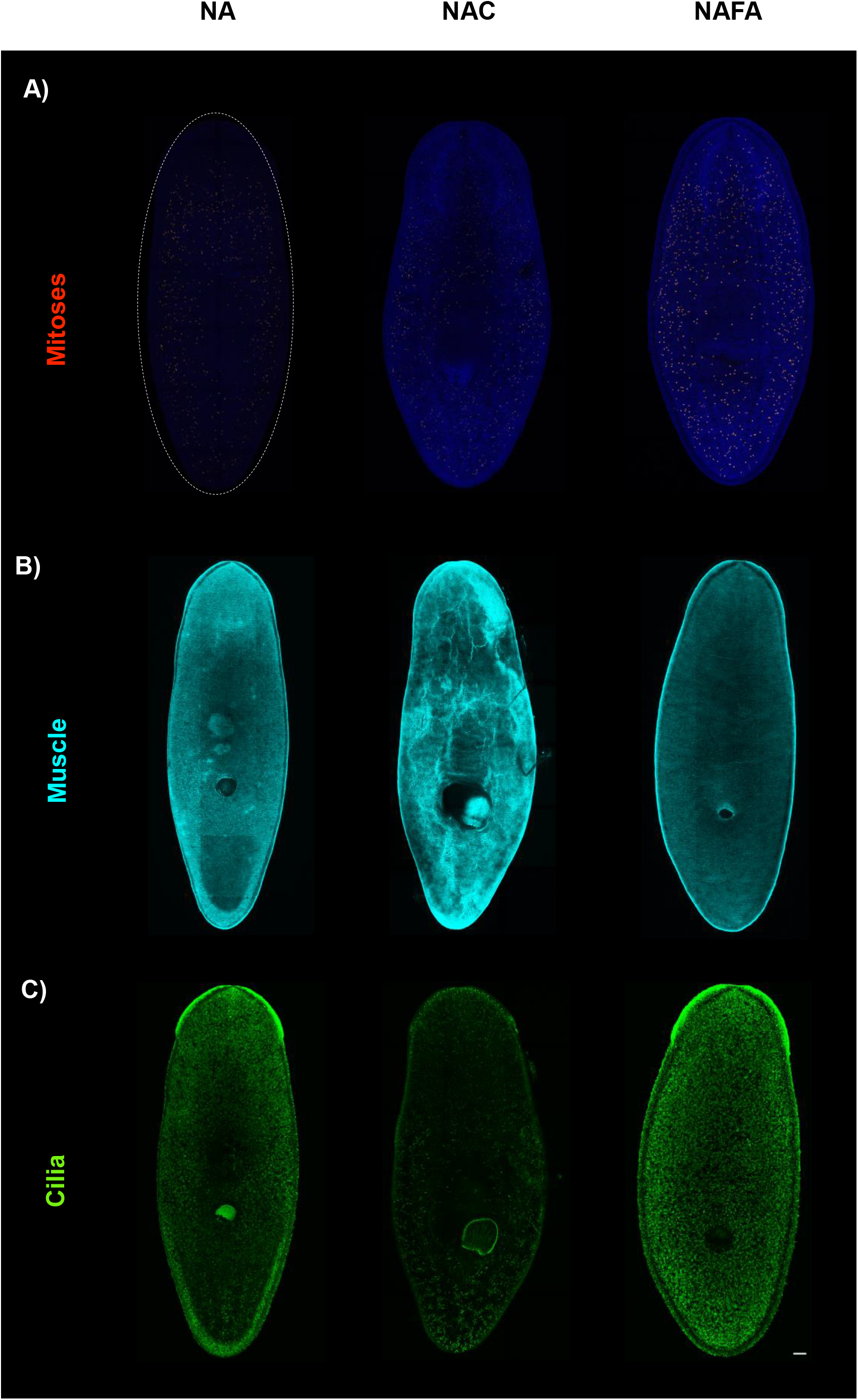
NAFA protocol enables immunofluorescent detection after fluorescent *in situ*. Immunostaining of **(A)** anti-phosphorylated Histone H3 (mitoses), **(B)** 6G10-2C7 (muscle) and **(C)** anti-acetylated tubulin (cilia) antibodies. Maximum intensity projection of confocal images, scale bar: 100 μm

We then assessed if we could use the new NAFA protocol to develop two-color FISH with two different RNA probes. *piwi-1* and *zpuf-6* were selected as targets for these double FISH experiments. Because the NA protocol is not compatible with ISH, we only compared the NAFA and NAC protocols to each other. First, we detected *zpuf-6* gene expression followed by *piwi-1*. We used confocal microscopy to image the samples, and observed similar expression patterns of *piwi-1* in both protocols. However, the NAFA protocol showed a clearer expression pattern of the epidermal progenitor *zpuf-6* because the integrity of the epidermis was preserved (Figure 5A and 5B). After the double FISH we explored the mitotic cells in the same samples using anti-H3P antibody. The immunostaining again showed stronger signal for mitotic cells with the NAFA protocol (Figure 4A, Figure 5A, and Figure 5B). Therefore, NAFA showed the greatest improvements in FISH and immunostaining development without affecting the planarian epidermis.

**Fig 5.**
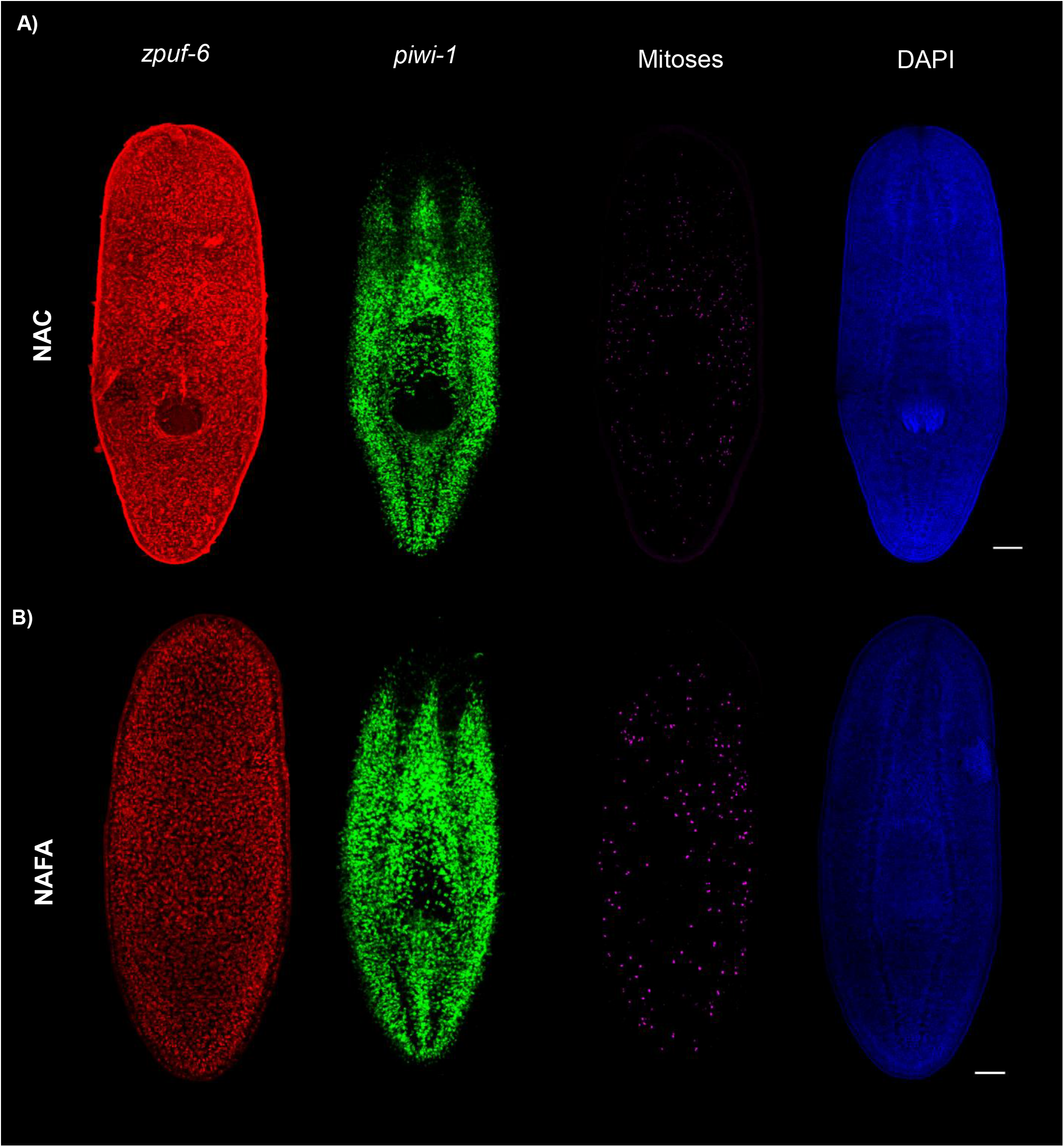
NAFA protocol is compatible with double fluorescent *in situ* hybridization followed by immunofluorescence. **(A and B)** FISH of *zpuf-6* and *piwi-1*. **(A and B)** Immunostaining of phosphorylated Histone H3 (mitoses). Maximum intensity projection of confocal images, scale bars: 100 μm

To confirm that the NAFA protocol does not affect the epidermis after double FISH of *piwi-1* and *zpuf-6*, we performed immunostaining for cilia (Figure 6). The confocal images of the dorsal and ventral sides of planarians after two-color FISH showed well preserved cilia with the NAFA protocol. In contrast, we failed to detect the same pattern of cilia in planarians treated with NAC protocol (Figure 6A and 6B). Additionally, confocal images of *zpuf-6* expression showed that epidermal integrity was disrupted by the NAC protocol, but preserved by the NAFA protocol (Figure 6A and 6B).

**Fig 6.**
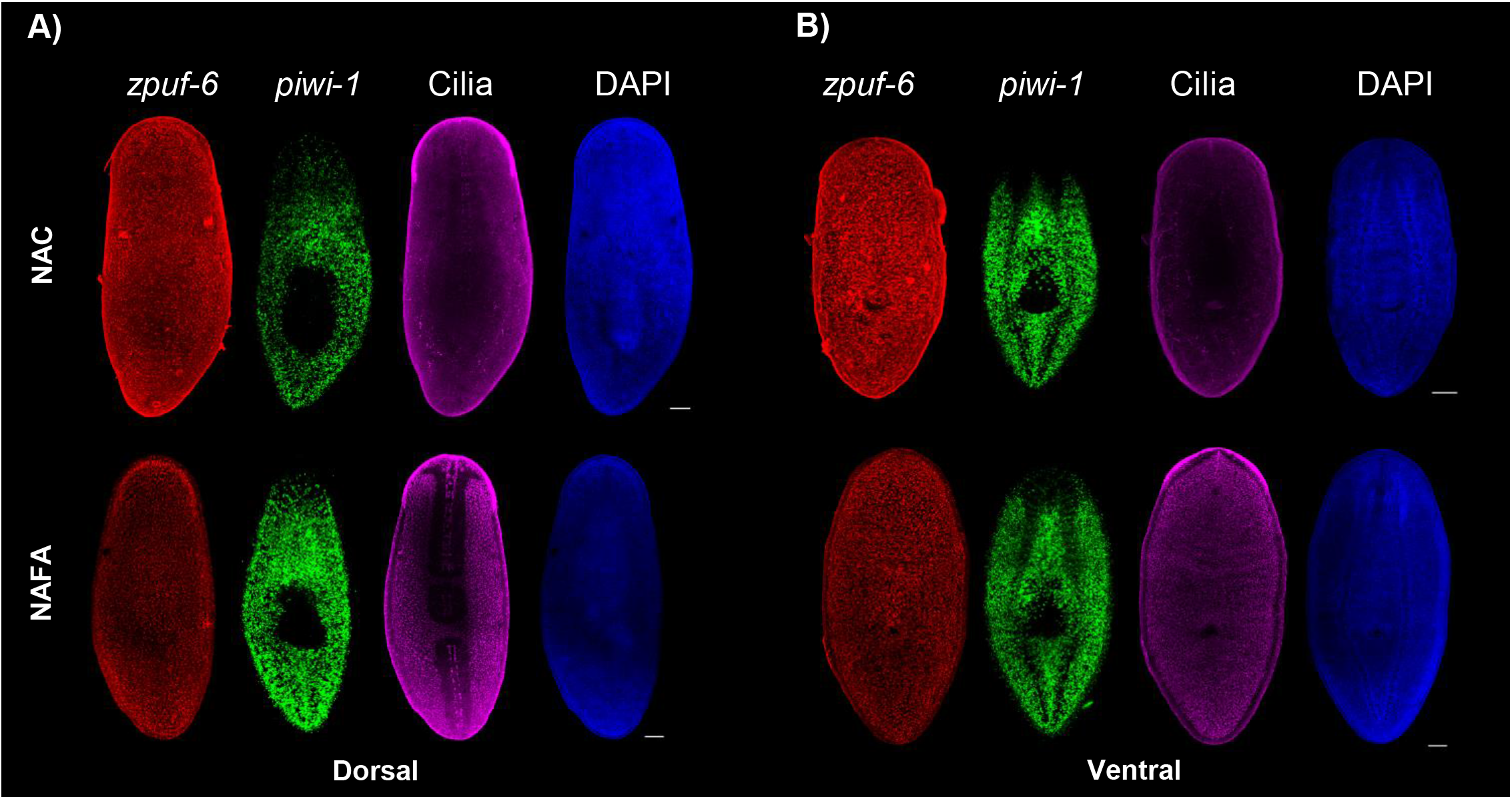
NAFA protocol maintains cilia on the epidermis after double fluorescent *in situ* hybridization. **(A and B)** FISH of *zpuf-6* and *piwi-1* and immunostaining of antiacetylated tubulin (cilia). Maximum intensity projection of confocal images, scale bars: 100 μm

We next tested if we could use the NAFA protocol to study the wounding response during planarian regeneration without affecting the epidermal or blastema tissue. We performed FISH of *piwi-1* and the immunostaining of cilia on trunk fragments after 8 hpa (hours post amputation), 1 dpa, 2 dpa, 4 dpa and 8 dpa (days post amputation) to assay for epidermal integrity (Figure 7A and 7B). Confocal images showed that epidermal integrity was not maintained when the NAC protocol was used. In contrast, we could see very clear staining of cilia on trunks during regeneration using the NAFA protocol (Figure 6B, 7A and 7B). Likewise, the anterior and posterior blastema of trunk fragments treated with the NAFA protocol remained intact (red arrows on Figure 7B). Remarkably, while the *piwi-1* FISH pattern was similar between the NAFA and NAC protocols, the confocal images of the different fragments during regeneration revealed an area of undifferentiated tissue that could not be detected when using the NAC protocol (Figures 6 and 7). Finally, to further investigate blastema integrity and ISH signal, we performed FISH in tandem with immunostaining. High magnification confocal imaging showed that the NAFA protocol preserved the wound epidermis and the blastema, while it was heavily damaged by the NAC protocol (Figure 7C and S3). In conclusion, the NAFA protocol preserves the delicate wound epidermis and blastema during regeneration.

**Fig 7.**
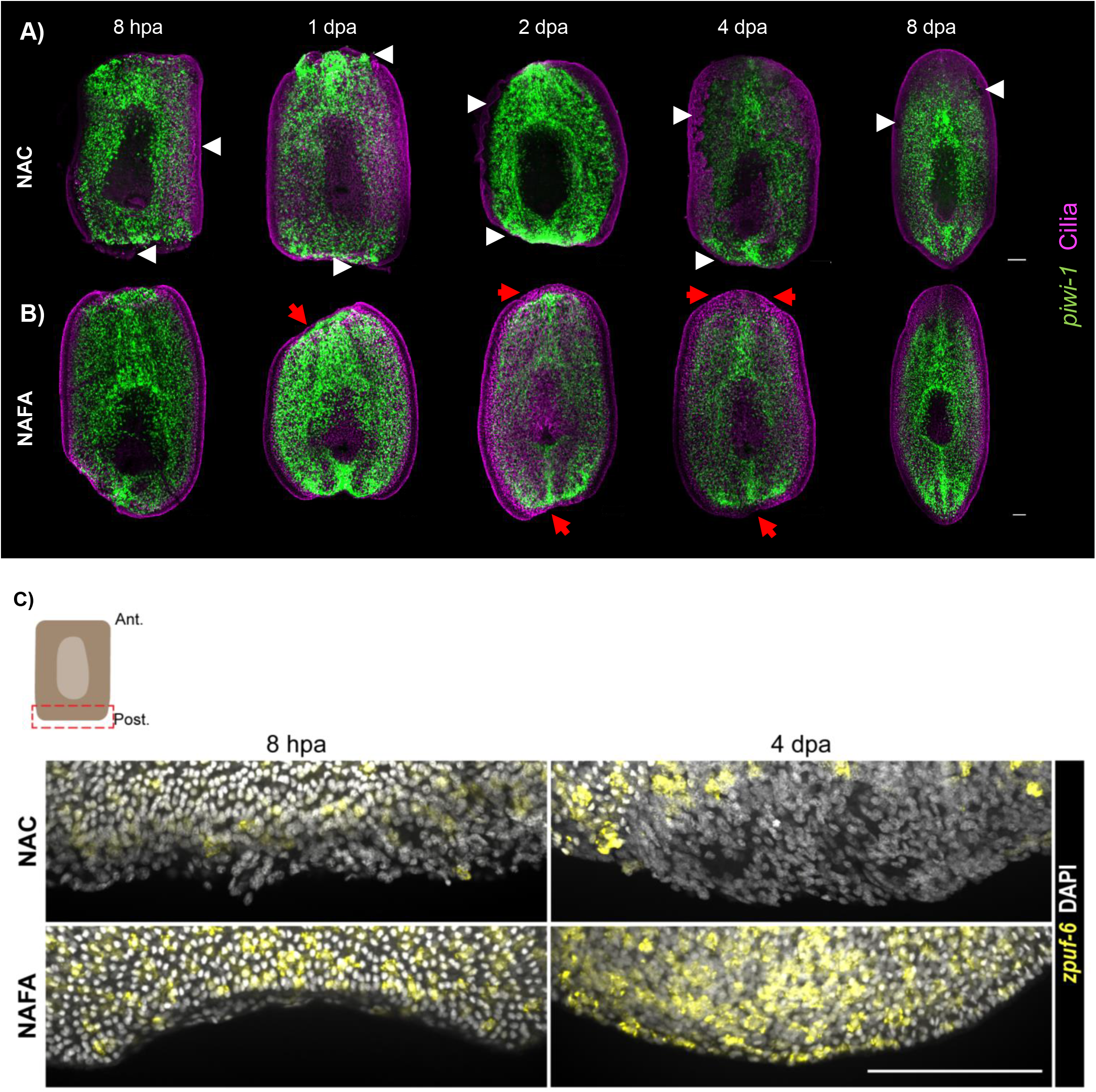
NAFA protocol maintains epidermal integrity during regeneration. **(A and B)** FISH of *piwi-1* and immunostaining of anti-acetylated tubulin (cilia). **(A)** White arrows show the affected epidermal layer and **(B)** red arrows at the blastema show epidermal integrity during regeneration. **(C)** FISH of *zpuf-6* and DNA staining with DAPI. Maximum intensity projection of confocal images, scale bars: 100 μm

## Discussion

The new NAFA protocol reported here works well for both FISH and WISH in the planarian *S. mediterranea*. While existing protocols work well, theu are only suitable for either immunohistology or ISH. However available protocols do not work robustly when both FISH and WISH need to be applied to the same sample. Additionally, current ISH protocols include harsh treatments to facilitate probe penetration which damage the ciliated epidermis and blastema in these animals. Preservation of external tissue layers is especially important for a research organism used in the study of regeneration, as stem cell proliferation and differentiation take place just beneath the wounding epidermis to form a blastema which grows to replace lost tissues^2^. The new NAFA protocol addresses these shortcomings and allows for performing immunohistology and ISH on the same samples while preserving the delicate outer cellular layers of this organism. The marked tissue preservation provided by the fixation protocol we describe here will enable researchers to apply FISH and WISH to the same sample thus opening the door to a myriad of new experiments.

The NAFA protocol, like the NA protocol, is highly compatible with immunofluorescence. Both protocols use nitric acid during fixation, which is known to euthanize and flatten planarians while preserving the ciliated epidermis ^13, 14, 25^. However, use of nitric acid alone is not sufficient to detect gene expression using ISH. To develop a protocol that is compatible with both ISH and immunofluorescence, we explored the use of carboxylic acids, which are widely used in a variety of fixation approaches^26^. These methods are a subset of a broader class called coagulant fixatives which act by precipitating proteins instead of covalently crosslinking them^16^. Acid treatments enhance immunohistochemical studies by hydrolyzing crosslinks and potentially disrupting protein complexes, in a process known as antigen retrieval^27^. In contrast, the NAC protocol uses enzymatic proteinase K treatment to permeabilize the sample. While weak immunofluorescence signals can be generated from this method, these signals are much weaker than those produced by the NAFA or NA methods, presumably due to the loss of target epitopes by enzymatic digestion. Furthermore, the harsh mucolytic NAC treatment tears the outer layers of the planarian body, making it difficult to use for studying fragile tissues like the epidermis or regeneration blastema.

The NAFA protocol is also highly compatible with *in situ* hybridization. This is in stark contrast to the NA protocol. Three main possibilities exist to account for this compatibility: 1) that samples fixed using the NAFA protocol are more permeable to riboprobes than samples fixed by the NA protocol are; 2) that RNA targets are more available to ISH probes than they are in other coagulating fixation conditions; or 3) that target RNA molecules are more intact than they are in samples that have harsher acid treatments. We evaluate the likelihood of each of these three possibilities.

First, samples fixed with the NA protocol are sufficiently permeabilized to allow antibodies to penetrate to internal structures detectable by immunofluorescence, yet *in situ* hybridization fails on these samples. While the structures of specific antisense mRNA probes are unknown, the relatively short probes used in this manuscript still do not yield any appreciable signal with the NA protocol. Therefore, sample permeability may not explain NAFA’s superior performance in ISH. Since the size of riboprobe affects its diffusion rate and penetration, a systematic study with riboprobes of varying lengths is necessary to assess permeabilization in samples fixed by both the NA and the NAFA protocols. Second, relative to prolonged strong acid treatments, such as the NA protocol, the proteins in NAFA samples will likely not be hydrolyzed to the same extent, and will also be crosslinked, two factors which would be expected to increase the size and complexity of proteins bound to and around RNA molecules. Since NAFA fixation likely leads target RNA molecules to be bound or surrounded by networks of crosslinked proteins, we believe increased RNA availability to probes is another unlikely explanation for the compatibility of NAFA with ISH. Third, compared to the NA protocol, NAFA’s much briefer nitric acid treatment almost certainly results in less acid hydrolysis of RNA. Furthermore, the NAFA protocol includes EGTA to chelate calcium and magnesium ions, as many RNase enzymes require these to digest RNA molecules^17^. Of the three possibilities for the NAFA protocol’s compatibility with ISH, we posit that preservation of RNA integrity is the most likely explanation.

The benefits of the NAFA protocol are likely due to the unique approach of simultaneously performing crosslinking and carboxylic acid treatments. As we devised this method, we tested three carboxylic acids for their performance in ISH and chose formic acid, which is chemically the smallest and simplest carboxylic acid, for use in the NAFA protocol. Formic acid is the strongest of the three acids tested in this manuscript. It is unknown whether other untested carboxylic acids would perform better on ISH in planarians. However, for aliphatic carboxylic acids such as the ones tested here, increasing length of the carbon chain is inversely proportional to acid strength, so we expect other acids would be unlikely to produce the full benefits created by the formic acid treatment of the NAFA protocol. Furthermore, carboxylic acids with long aliphatic carbon chains have detergent-like properties, making them potentially unsuitable for fixing tissue samples.

The NAFA protocol may have uses beyond preparing whole-mount planarian samples for IF or ISH. Because it preserves the integrity of the ciliated epidermis in planarians, this method may be useful for the study of other samples with multiciliated cells. Examples of multiciliated tissues include the lung epithelium, oviduct, and inner ear. This protocol may also be useful for the study of other animals, particularly highly regenerative animals with fragile regeneration blastemas. Future work will explore the applicability of the NAFA protocol in a diverse array of samples and research organisms.

## Acknowledgments

We thank Yongfu Wang and Nancy Thomas for assistance with histological procedure and all the people in the Histology core. We thank Cindy Maddera and Sean McKinney for assistance with confocal microscopy. We thank the Jerry Workman laboratory and the Proteomics Core for generously sharing reagents. We thank members of the Sánchez Alvarado laboratory for useful feedback. We thank Hanh Vu for her generous help and technical assistance.

## Supplementary figures

**Supplementary Fig 1.**
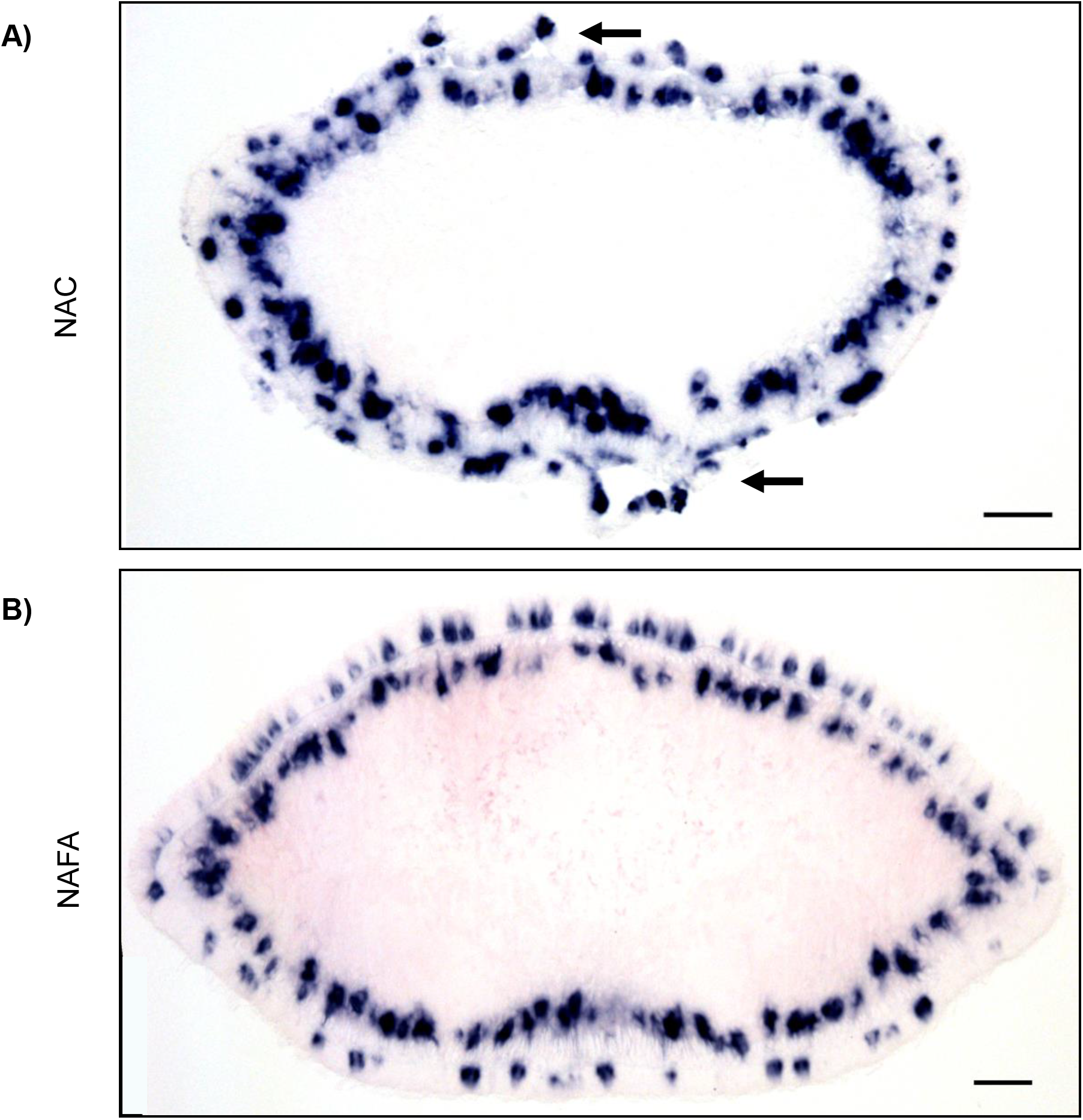
Epidermal integrity is preserved with NAFA protocol. Chromogenic *in situ* of *zpuf-6*. Anterior transverse histology section. (A) Black arrows show the affected epidermal layer. Scale bars: 100 μm

**Supplementary Fig 2.**
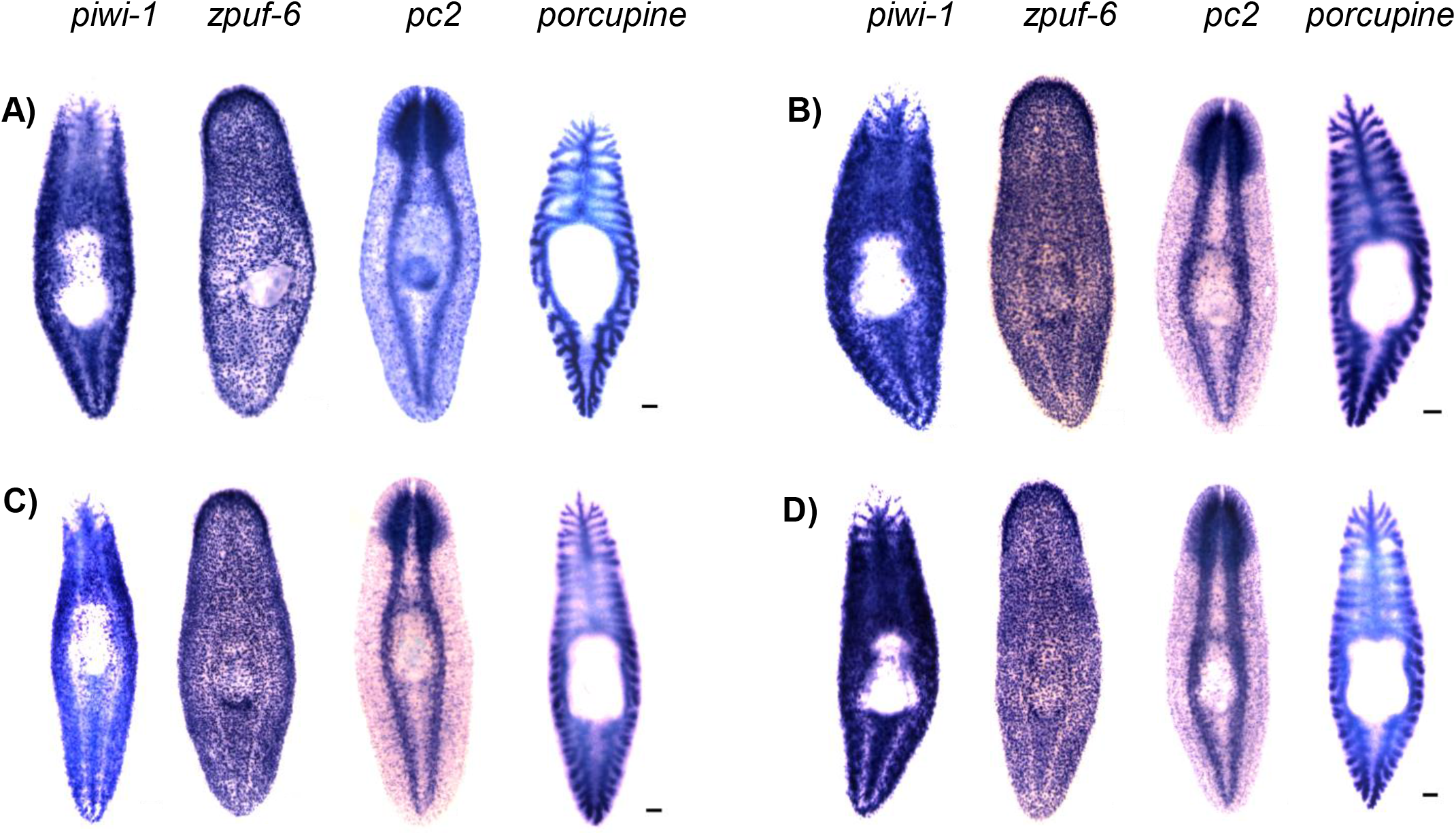
Different carboxylic acid tested in optimization of a new *in situ* protocol. Chromogenic WISH of *piwi-1, zpuf-6, pc2*, and *porcupine*. **(A)** NAC, **(B)** formic acid (4.8%), **(C)** acetic acid (4.9%) and **(D)** lactic acid (4.2%). Scale bars: 100 μm

**Supplementary Figure 3.**
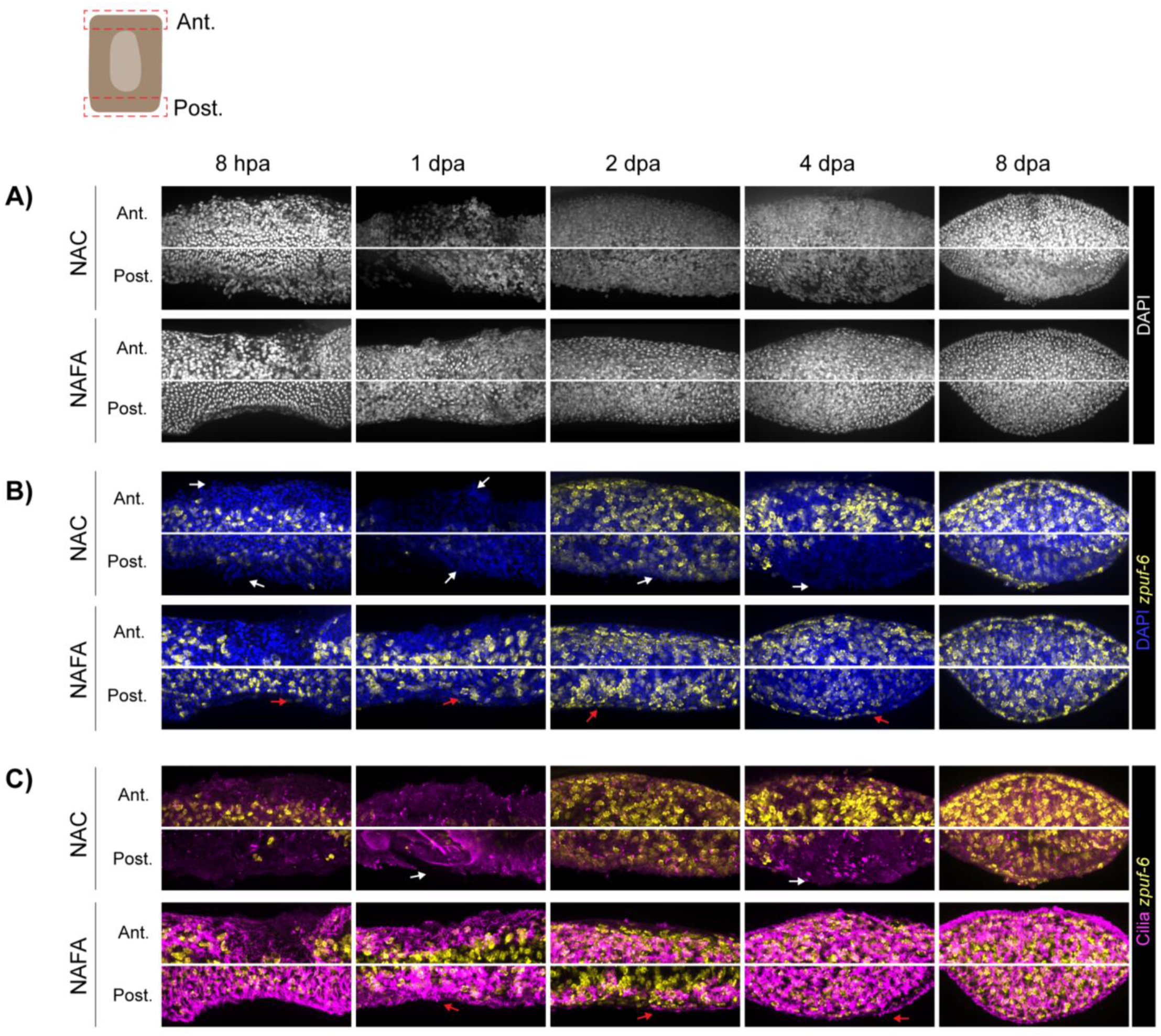
NAFA protocol maintains epidermal and blastema integrity during regeneration. **(A)** DAPI staining. **(B)** FISH of *zpuf-6* and DAPI staining. **(C)** FISH of *zpuf-6* and immunostaining of anti-acetylated tubulin (cilia). **(B and C)** White arrows show the affected epidermal layer and red arrows at the blastema show epidermal integrity during regeneration. Maximum intensity projection of confocal images (40X). Ant: anterior and Post: posterior. Regenerating trunk fragments.

